# Genomic selection validated across two generations of loblolly pine breeding

**DOI:** 10.64898/2026.01.22.701135

**Authors:** Fikret Isik, Mohammad Nasir Shalizi, Trevor Walker

## Abstract

This study evaluated the effectiveness of genomic selection (GS) in loblolly pine (*Pinus taeda*) using a two-generation closed breeding population and a genetically diverse Mainline population. Single-step genomic best linear unbiased prediction (ssGBLUP) models were used to include all phenotypic, genotypic, and pedigree information. Prediction accuracies of genomic estimated breeding values reached up to 0.70 for stem volume and stem straightness. Prediction accuracy showed a strong linear relationship with mean relatedness between training and validation populations (r > 0.92). Adjusting the scaling between genomic and pedigree relationship matrices improved model stability, increased prediction accuracy, and reduced bias in genomic estimated breeding values. Estimates of heritability and variance components from ssGBLUP were consistent with pedigree-based models, particularly when genomic relationships were properly scaled. Genomic selection had approximately 50% more genetic gain per year relative to conventional selection. Overall, these results demonstrate that GS can be effectively integrated into operational conifer breeding programs, given sustained investment in large, well-connected training populations with high-quality phenotypic data. We also outline the planned implementation of GS in the North Carolina State University Cooperative Tree Improvement Program to increase genetic gain.

## Introduction

Genomic selection (GS) refers to predicting breeding values for individuals without phenotypic records using genome-wide DNA markers. Unlike early marker–trait association approaches that focused on a small number of loci with large effects, GS embraces the polygenic architecture of quantitative traits by modeling the covariance among relatives rather than individual marker effects. Since its introduction (Meuwissen et al. 2001), GS has triggered a paradigm shift in plant and animal breeding. In U.S. dairy cattle, for example, widespread adoption of GS has doubled the rate of genetic gain since 2010 (Wiggans and Carrillo 2022). GS has also been successfully integrated into major crop improvement programs, including maize, wheat, and rice (Dreisigacker et al. 2021; da Silva et al. 2025).

For perennial, outcrossing woody species like forest trees, long generation intervals substantially slow genetic improvement when relying on conventional progeny testing (Dreisigacker et al. 2021; da Silva et al. 2025). By reducing the need for progeny testing, GS can accelerate breeding cycles and increase gain per unit time. Early reviews highlighted the promise of GS for forest trees (Grattapaglia 2014; Isik 2014), and numerous proof-of-concept studies using intragenerational cross-validation methods demonstrated encouraging results (e.g., Resende et al. 2012; Ratcliffe et al. 2015; Durán et al. 2017; Shalizi et al. 2022; Lauer et al. 2022). These early studies partitioned a single generation into training and prediction sets to mimic the application of GS, which can overestimate prediction accuracy because marker–QTL linkage phases are largely preserved between subsets (Grattapaglia 2022; Isik 2022).

A more realistic assessment of GS performance uses earlier generations as the training population and their descendants as the validation population (Bartholomé et al. 2016). This approach accounts for the effects of recombination, which disrupts marker-QTL associations across generations, and therefore better reflects operational GS performance in true breeding programs. Despite growing interest in GS for forest trees, true across-generation validation remains rare, with only a few recent examples (Simiqueli et al. 2023; Duarte et al. 2024). Reported accuracies vary widely and are influenced by training population size, phenotypic data quality, and the degree of relatedness between training and validation sets. Among these factors, relatedness plays a critical role in GS prediction accuracy but has not been thoroughly quantified in operational breeding contexts (Isik 2022). Clear guidelines are therefore needed to define appropriate training populations and to understand the limitations of GS when applied across generations in species with long breeding cycles.

Historically, GS was implemented using multi-stage genetic evaluation pipelines (VanRaden et al. 2009). In these approaches, pedigree-based best linear unbiased prediction (ABLUP) was first used to estimate breeding values, which were then converted into pseudo-phenotypes for genomic prediction (GBLUP). Genomic breeding values for genotyped individuals were subsequently combined with pedigree-based estimates using selection indices (Christensen et al. 2012; Bermann et al. 2022). These multi-stage approaches can introduce bias, reduce prediction accuracy, and do not fully utilize genomic information for related but non-genotyped individuals (Misztal et al. 2009).

To address these limitations, single-step genomic best linear unbiased prediction (ssGBLUP) was proposed (Legarra et al. 2009; Christensen and Lund 2010), which integrates phenotypic, pedigree, and genotypic information into a unified framework (Lourenco et al. 2020). Compared with conventional ABLUP or two-stage GBLUP approaches, ssGBLUP is simpler to implement, reduces bias between genotyped and non-genotyped individuals, accounts for selection on Mendelian sampling within families, and improves prediction accuracy (Misztal and Legarra 2017; Lourenco et al. 2020; Mäntysaari et al. 2020). Advances in computational efficiency, including sparse matrix algorithms and faster construction of the inverse hybrid relationship matrix, have enabled routine application of ssGBLUP to large-scale breeding datasets, making it a standard method in modern plant and animal breeding programs (Bermann et al. 2022; Misztal et al. 2021). The value of ssGBLUP has been demonstrated for in forest tree breeding (Walker et al. 2022; Ukrainetz and Mansfield 2020; Gonzalez et al. 2025), but its implementation in operational tree breeding programs has yet to be realized.

For ssGBLUP to perform optimally, genomic relationships derived from SNP markers must be compatible with pedigree-based relationships for genotyped individuals (Legarra et al. 2014). Incompatibility arises primarily because allele frequencies of the base population, a theoretical group of unrelated individuals that serve as founders, are unknown and are typically approximated using the genotyped population’s allele frequencies (Vitezica et al. 2011; Christensen 2012). This can lead to biased estimates of variance components and breeding values (Forni et al. 2011). Several approaches have been proposed to make the relationships compatible, including regressing genomic relationships toward pedigree relationships (VanRaden 2008), adjusting the genomic relationship matrix by adding the average difference between pedigree and genomic relationships (Vitezica et al. 2011), and introducing meta-founders to explicitly model base population relationships, analogous to genetic groups in animal models (Westell et al. 1988; Christensen et al. 2012; Legarra et al. 2015). The evolution of these methods and their implications for ssGBLUP have been comprehensively reviewed by Bermann et al. (Bermann et al. 2022). However, their application in tree breeding remains limited.

In this study, we evaluated GS using a multi-generational population of loblolly pine (*Pinus taeda*) developed by the Cooperative Tree Improvement Program at North Carolina State University. This closed breeding population, founded with 21 parents, was bred over two generations (2001–2022), and has pedigree connections with a larger, more diverse mainline population. The objectives of the study were to:

1. Validate GS in a multi-generational breeding population by comparing its efficiency with conventional selection in terms of genetic gain per unit time.
2. Assess how relatedness between training and validation sets and training set size influence prediction accuracy using a large operational breeding population; and
3. Evaluate the effect of different weights assigned to pedigree- and marker-based relationships in ssGBLUP to address incompatibility between relationship matrices.

## Materials and Methods

### Plant Material

#### First generation Atlantic Coastal Elite population (ACE1)

The first-generation Atlantic Coastal Elite (ACE1) population was developed from three disconnected eight-parent diallels (Shalizi and Isik 2019). In total, 24 founding parents were mated to produce 76 full-sib families. Seedlings were artificially inoculated in 2007 with a mixed inoculum of *Cronartium quercuum* f. sp. *fusiforme*, the causal agent of fusiform rust disease. Individuals that developed stem galls were culled, and the remaining 2488 gall-free seedlings, representing 51 full-sib families from 21 parents and 7 pollen-mix families were clonally propagated via rooted cuttings.

Eight field trials were established in 2009 and 2010 using an incomplete row–column design (**Figure 1A**). Each clone was represented by a single copy (ramet) per site. Additionally, 443 seedlings from a checklot family were included across trials, bringing the total number of planted trees to 14496 (**Table 1**). The ACE1 field trials were assessed in 2016, six years after establishment.

**Figure 1.**
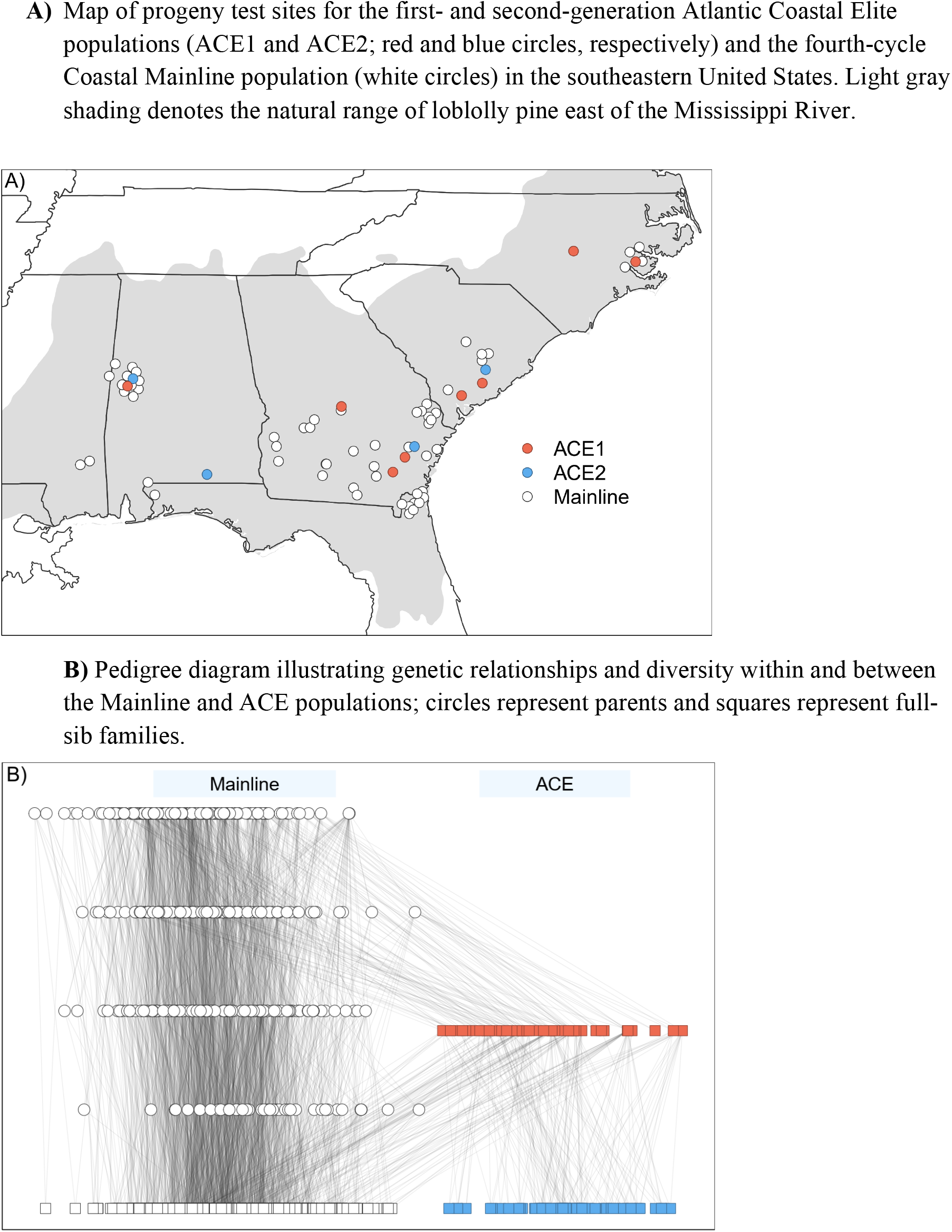
Location and pedigree of the study population. **A)** Map of progeny test sites for the first- and second-generation Atlantic Coastal Elite populations (ACE1 and ACE2; red and blue circles, respectively) and the fourth-cycle Coastal Mainline population (white circles) in the southeastern United States. Light gray shading denotes the natural range of loblolly pine east of the Mississippi River. **B)** Pedigree diagram illustrating genetic relationships and diversity within and between the Mainline and ACE populations; circles represent parents and squares represent full-sib families.

**Table 1.** Summary of population structure for the Atlantic Coastal Elite (ACE1 and ACE2) and Coastal Mainline populations of loblolly pine (*Pinus taeda*), including the number of field tests, total and unique trees, parents, full-sib crosses, and the distribution of genotyped and non-genotyped individuals used for genetic evaluation.

|  | **ACE1** | **ACE2** | **Mainline** |
| --- | --- | --- | --- |
| Field tests | 8 | 4 | 60 |
| Trees | 14496 | 3639 | 47991 |
| Unique trees | 2931 | 3639 | 47991 |
| Parents | 28 | 85 | 497 |
| Crosses | 51 | 73 | 813 |
| Genotyped trees | 2076 | 2108 | 6140 |
| Non-genotyped trees | 855 | 1531 | 41851 |

#### Second generation Atlantic Coastal Elite population (ACE2)

In 2017, 73 ACE1 clones were selected with emphasis on growth rate and stem form and mated to produce 67 full-sibling families for progeny testing (ACE2 generation). To enhance genetic connectivity with the Mainline population, the ACE2 field trials also included 6 full-sib crosses among ACE1 clones and parents from the Mainline population, as well as the same 7 pollen-mix families used in the previous ACE1 generation. Approximately 3639 seedling progeny from these families were planted at four locations in 2021 using incomplete row–column designs (**Table 1, Figure 1A**).

#### Mainline Coastal breeding population

The Atlantic Coastal Mainline fourth-cycle breeding population consists of about 497 parents. These parents were mated between 2011-2019 to produce 813 full-sib families (Isik and McKeand 2019). Progeny tests were established at 60 sites between 2014 and 2022, each containing about 120 families replicated in six to twenty blocks using incomplete row–column designs (**Table 1, Figure 1A**). On average, a full-sib family had around 60 progeny at planting. The Mainline population shares pedigree connections with the ACE populations described before, including 28 and 40 parents shared with ACE1 and ACE2 populations, respectively, as well as common ancestors (**Figure 1B**).

#### Phenotyping

The ACE1 trials were assessed at age six years and the ACE2 and Mainline population trials were assessed at age four years. Measurements included individual-tree height (m), diameter at breast height (cm), stem straightness (1 to 6, with 1 indicating the straightest stems), incidence of fusiform rust disease (yes or no), and occurrence of stem forking (yes or no). Outside bark stem volume (dm^3^) was calculated using the volume equations (Sherrill et al. 2008). Approximately, 14496 trees were evaluated in the ACE1 trials, 3639 in the ACE2 trials, and 47991 in the Mainline trials. In this study, stem volume and stem straightness were analyzed as the response variables.

#### Genotyping

In total, 10324 trees from the ACE1, ACE2, and Mainline populations were genotyped, along with 213 founder and ancestral trees (**Table 1**). Diploid needle tissue samples were collected from the fresh growth of trees. Needle samples were placed in labeled coin envelopes and stored with silica desiccant to preserve freshness. Once dried, approximately 10 mg of tissue was prepared for DNA extraction. Trees were genotyped using the Pita50K Axiom™ SNP array developed for loblolly pine (Caballero et al. 2021). SNP markers were filtered based on a minor allele frequency threshold of 0.01 and a maximum missing rate of 10%. Samples with a no-call rate exceeding than 10% were excluded from further analysis. After filtering, approximately 15000 SNP markers were retained with ∼9% of genotypes call missing. Missing genotypes were imputed using the “wright” method of the *snpReady* package (Granato et al. 2018) in R statistical environment (R Core Team 2024).

### Statistical Models

Pedigree-based linear mixed models (ABLUP) were fitted to estimate variance components and obtain empirical breeding values (EBVs). Single-step genomic best linear unbiased prediction (ssGBLUP) models were fitted using the same fixed and random effects to estimate variance components and genomic estimated breeding values (GEBVs).

#### Predictions of empirical breeding values using ABLUP

The following individual-tree (animal) model was fit to predict empirical breeding values (EBVs) on the same scale across generations:

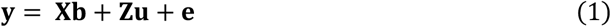

where **y** is the vector of observations; **X** and **Z** are incidence matrices for fixed and random effects, respectively; **b** is the vector of fixed effects (intercept and site effects); **u** is the vector of random effects (including additive and dominance genetic effects and experimental designs factors) and **e** is the vector of residuals. The vector of observations was assumed to follow a multivariate normal distribution with following expectations:

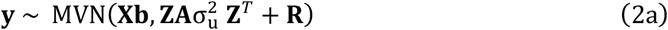

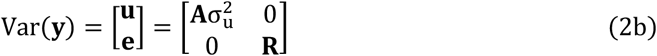

where **A** is the pedigree-based relationship matrix, 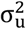 is the additive genetic variance explained by breeding values, and **R** is the residual covariance matrix. Random specific combining ability effect (cross effect), experimental design factors replication, row, and column effects were assumed to have multivariate normal distribution with expectations of ∼*NID*(0, *σ*^2^). They were not included in equation 2 for simplicity. Similarly, in the baseline model, residuals were assumed to have the expectations of 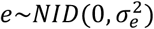. Where appropriate, these assumptions were relaxed by fitting block-diagonal covariance structures within each test site, expressed as direct sums of the form 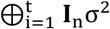 to allow heterogeneous variance structures (Isik et al. 2017). The symbol ⊕ denotes the direct sum operator, and **I**_n_ is an identity matrix of appropriate dimensions. Henderson’s (1982) mixed model equations were solved to estimate EBVs:

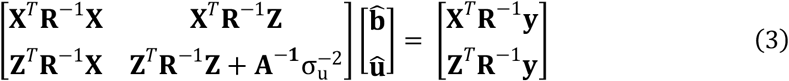

Where 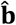 is the vector of best unbiased linear estimates of fixed effects and 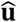 is the vector of best linear unbiased predictions of random effects, 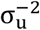 is the inversed additive genetic variance, and **R**^−1^ is the inversed residual covariance matrix. Again, for brevity, additive genetic effects (tree) were included in mixed model equations only.

#### Predictions of genomic breeding values using ssGBLUP

In this study, only a subset of the individuals was genotyped due to the cost of marker assays and logistical constraints (**Table 1**). To fully exploit all available phenotypic, pedigree, and genomic information, we applied single-step genomic BLUP (ssGBLUP), in a single linear mixed-model framework. The linear mixed model and assumptions were identical to those described in Equations (1) and (2), except that the additive genetic effects (**u**) were modeled using a hybrid relationship matrix (**H**), which integrates pedigree-based (**A**) and marker-based (**G**) genetic relationships matrices. In practice, the inverse of **H** is used because it is computationally more efficient to construct than **H** itself (Aguilar et al. 2010; Christensen and Lund 2010).

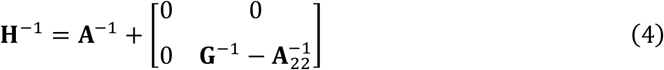

where **A**^−1^ is the inverse of the full **A** matrix, 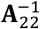 is the inverse of the pedigree-based submatrix for genotyped individuals, and **G**^−1^is the inverse of the genomic relationship matrix for the same individuals. Additive genetic relationships based on pedigree are subtracted from marker-based relationships of genotyped individuals to prevent double counting.

#### Compatibility of genomic and pedigree-based relationships

The genomic relationships matrix (**G)** among genotyped individuals was estimated using the allele frequency method of VanRaden (2008). Let **M** be the matrix of gene content centered on 0, where each element takes the value −1, 0, or 1, corresponding to homozygous for the major allele, heterozygous, and homozygous for the minor allele, respectively. Marker genotypes were centered by subtracting twice the allele frequency at each locus (2*p*_*i*_). The **G** matrix was computed as

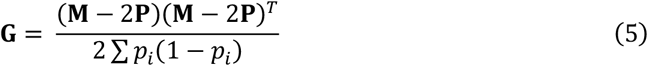

where *p*_*i*_ is the minor allele frequency at marker *i*, **P** is a vector of allele frequencies, 2 ∑ *p*_*i*_(1 − *p*_*i*_) represents the total marker-based additive genetic variance summed across all *m* loci.

For ssGBLUP, **G** must be compatible with the pedigree-based relationship matrix among genotyped individuals (**A**_22_), which assumes a base population of founders having mean relationships of zero. To address incompatibility, first we aligned the **G** matrix with the **A**_22_ matrix using the regression method according to Christensen et al. (2012);

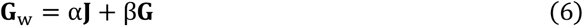

where **J** is the unity matrix, α and β are the intercept and the slope, respectively. The regression parameters were estimated by equating the elements of two matrices, **G** and **A**_22_ and the means of their diagonal elements as follows (Meyer et al. 2018);

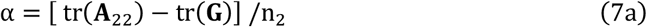

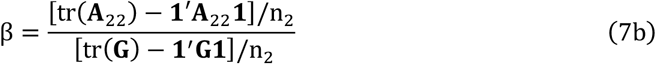

where tr(·) denotes the trace (sum of diagonal elements), n is the number of genotyped individuals and **1′** is the transposed vector of unity. This scaling ensures that **G** and **A**_22_ are on the same genetic base, enabling unbiased integration into the ssGBLUP model (Bermann et al. 2022). Then, a scalar weighting parameter λ was applied to scale the genomic and pedigree relationships to reduce the bias in predictions of GEBV (Aguilar et al. 2010).

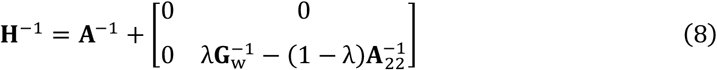

The parameter λ ranges from 0 and 1, where λ = 1 assigns full weight on genomic relationships for genotyped trees, whereas smaller values of λ shrink genomic relationships toward the pedigree relationships, thereby increasing the contribution of **A**_22_. Reducing λ can be advantageous for traits of lower heritability, or when SNP markers capture only a portion of the additive genetic variance (Meyer et al. 2018). We evaluated λ values of 0.9, 0.8, 0.7, 0.6, and 0.5 to assess the sensitivity of variance component estimates and the predictions of GEBV to the relative weighting of genomic information in **H**^−1^. For stem volume, a value of λ = 0.50 was used in the final analysis, as equal weighting of marker- and pedigree-based relationships resulted in improved model performance. For stem straightness, the default value of λ = 0.90 was retained in the final analysis.

### Genomic Selection Validation

#### Scenario 1: ACE1 as training and ACE2 as validation

In this scenario, EBVs for the two-generation ACE population were first estimated using the pedigree-based relationship matrix and phenotypes in the conventional ABLUP model. Next, the phenotypic records of the ACE2 population were masked and the ssGBLUP model was used to predict breeding values. The ssGBLUP model included phenotypes from ACE1 (2076 genotyped trees) as a training set but not for the ACE2 (2108 genotyped trees) as a validation set. The average genomic relationship between the training and validation sets was 0.036 (**Table S1**).

#### Scenario 2: ACE1 and Mainline as training and ACE2 as validation

In this scenario, the objective was to understand the effect of a larger training set on the prediction accuracy. We first fit the ABLUP model using combined phenotypic data from the two-generation ACE and Mainline populations to obtain EBVs. Next, the phenotypic records of the ACE2 population were masked and the ssGBLUP model was used to predict breeding values. This ssGBLUP model used ACE1 and Mainline as the training set (9071 genotyped trees) and ACE2 population as the validation set (2018 genotyped trees). The average genomic relationship between the training and validation sets was 0.024 (**Table S1**).

#### Scenario 3: ACE as training and Mainline population as validation

In this scenario, the two-generation ACE population served as the training set, while the Mainline population was designated as the validation set. First, EBVs were estimated for all phenotyped trees as in Scenario 2. Next, phenotypic records for 6140 genotyped trees in the Mainline population were masked. The ssGBLUP model was used to predict GEBVs using the genotyped Mainline trees as a validation set and the training population consisted of ACE trees. The average genomic relationship between the training and validation sets was 0.021 (**Table S1**).

#### The impact of genetic relatedness on prediction

The degree of genetic relatedness between the training and validation sets is a critical factor influencing the accuracy of GS (Shalizi et al. 2022; Lauer et al. 2022; Isik 2022). This is because SNP markers essentially capture genetic relationships more effectively than pedigree-based relationships (Forni et al. 2011). To evaluate this effect, the Mainline population was partitioned into validation subsets according to their average genomic relationship with the ACE training set. Twelve subsets were created, with average genomic relationships of 0.011 – 0.036 to the ACE population. Model validation statistics using the ACE as the training and the Mainline as validation were calculated separately for each subset to evaluate the impact of relatedness between the training and validation sets.

#### Validation metrics

We used linear regression analysis to evaluate the accuracy and potential bias of GEBVs in two validation populations (Legarra and Reverter 2018). In this analysis, EBVs of the validation set were regressed on their corresponding GEBVs, and several key validation metrics were derived:

- **Prediction accuracy**: calculated as the Pearson correlation between EBV and GEBVs. The EBVs are default selection unit in conventional methods, so we used them as the baseline.
- **Regression slope**: reflects prediction bias defined as the difference between means of true and predicted breeding values ū − û. This is the slope of regressing EBV on GEBV. A slope of 1 indicates unbiased prediction, values smaller than 1 suggest inflation (overestimation of GEBVs), while values greater than 1 suggest deflation (underestimation of GEBVs).
- **Root mean squared error (RMSE)**: quantifies the squared average prediction difference between EBV and GEBV and calculated here as 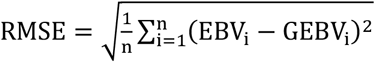.
- **Coefficient of determination (R**^**2**^): measures the proportion of variance in EBVs explained by GEBVs and obtained from linear regression.

## Results

### Genetic parameter estimates and response to selection

Genetic parameter estimates are reported from ABLUP and ssGBLUP models using a scaling factor (λ) of 0.50 for volume and 0.90 for straightness (**Table 2**). Genetic parameter estimates from the two models were similar and not statistically different. Individual-tree narrow-sense heritability for stem volume was low (0.11 to 0.13), whereas heritability for stem straightness was moderate (0.19 to 0.22). Additive genetic correlations between pairs of sites, reflecting the magnitude of genotype-by-environment interaction, were moderate for stem volume (0.70 for ABLUP and 0.66 and for ssGBLUP) and higher for stem straightness (0.83 and 0.87, respectively).

**Table 2.** Estimates of variance components, individual tree narrow-sense heritability (*h*^2^), and additive genetic correlation among sites (GxE) for stem volume and stem straightness in a two-generation Atlantic Coastal Elite (ACE) population of loblolly pine. Results are presented for pedigree-based BLUP (ABLUP) and single-step genomic BLUP (ssGBLUP) models. For ssGBLUP, marker- and pedigree-based relationship matrices were combined using scaling factor (*λ*) of 50% for stem volume and 90% for stem straightness. Estimates are followed by their standard errors (± SE).

| Parameters | Stem volume |  | Stem straightness |  |
| --- | --- | --- | --- | --- |
|  | ABLUP | ssGBLUP | ABLUP | ssGBLUP |
| Additive variance | 33 ± 2.7 | 30 ± 2.5 | 0.24 ± 0.015 | 0.29 ± 0.018 |
| Additive x site (GxE) | 15 ± 1.9 | 23 ± 2.7 | 0.03 ± 0.007 | 0.06 ± 0.011 |
| Residual variance | 216 ± 3.7 | 222 ± 3.6 | 0.96 ± 0.015 | 0.95 ± 0.015 |
| Heritability ($h^2$) | 0.13 ± 0.009 | 0.11 ± 0.009 | 0.19 ± 0.010 | 0.22 ± 0.012 |
| GxE correlation | 0.70 ± 0.032 | 0.66 ± 0.034 | 0.87 ± 0.023 | 0.83 ± 0.028 |

The distributions of breeding values (EBVs and GEBVs) for the first (ACE1) and second (ACE2) generations are shown in **Figure 2**. Genetic gain estimates for stem volume were 6.8% under both models (**Figure 2A**), whereas gains for stem straightness were 3.2% for EBVs and 2.6% for GEBVs (**Figure 2B**). Similar shifts in average EBV and GEBV across generations indicated that conventional selection and GS achieved comparable overall genetic gain across two ACE generations. The narrower and more peaked GEBV distributions observed in ACE2 indicate greater shrinkage relative to EBVs, apparently due to partial inflation of genomic predictions in the absence of phenotypes.

**Figure 2.**
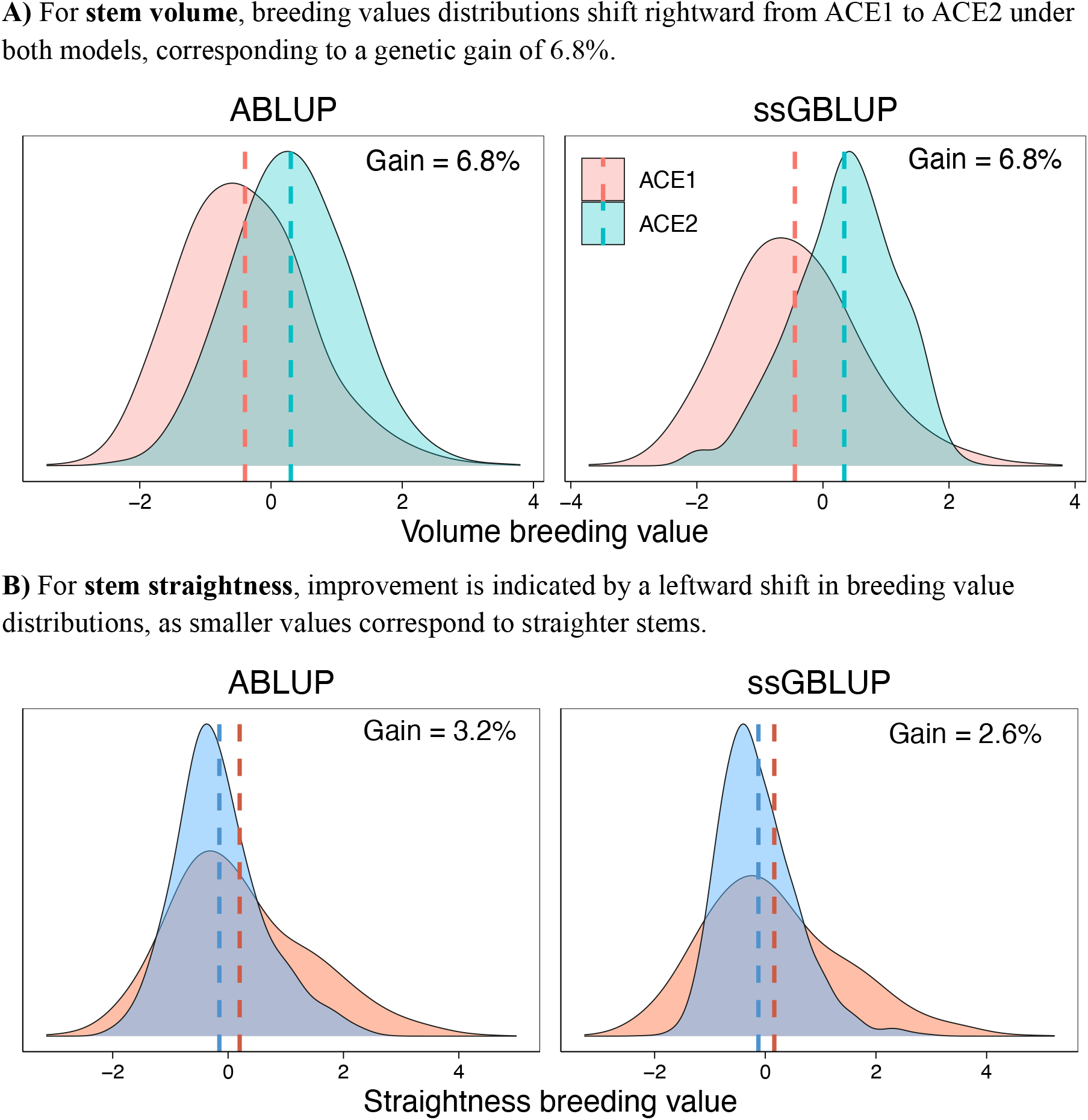
Distributions of breeding values across two generations of Atlantic Coastal Elite Population (ACE1 and ACE2) of loblolly pine, estimated using pedigree-based (ABLUP) and single-step genomic (ssGBLUP) models. Dashed vertical lines are population means. **A)** For **stem volume**, breeding values distributions shift rightward from ACE1 to ACE2 under both models, corresponding to a genetic gain of 6.8%. **B)** For **stem straightness**, improvement is indicated by a leftward shift in breeding value distributions, as smaller values correspond to straighter stems.

Assuming a breeding cycle length of 8 years for GS and 12 years for conventional selection, the annual rate of genetic gain was substantially higher using GS for stem volume (0.85% vs 0.57%) and for stem straightness (0.40% vs 0.21%). This represents an approximately 20%-50% increase in annual genetic gain attributable to cycle-time reduction.

### Validation of genomic selection

In the first validation scenario, the ACE1 served as the training population and ACE2 as the validation population. Prediction accuracy, measured as the correlation between EBVs and GEBVs, was moderate for stem volume (*r* = 0.55) and high for stem straightness (*r* = 0.67) (**Figure 3A**). Regression slopes for volume (*b*_1_ = 0.64) and stem straightness (*b*_1_ = 0.66) indicated inflation of GEBVs (**Table 3, Figure 3**), implying that the ssGBLUP model slightly overestimated the genetic merit of individuals at the lower tails of the distribution and underestimated those at the upper tail. Although this suggests moderate bias in the scale of GEBVs, their ability to rank individuals remains strong.

**Table 3.** Validation statistics for genomic prediction of stem volume and stem straightness in loblolly pine using the second-generation Atlantic Coastal Elite (ACE2) and Coastal Mainline populations. Model performance is reported for alternative training–validation combinations and summarized using the regression slope (bias), coefficient of determination (*R*^2^), and root mean squared error (RMSE). For ssGBLUP, marker- and pedigree-based relationship matrices were combined using scaling factor (*λ*) of 50% for stem volume and 90% for stem straightness.

| Training set | Validation set | Bias (slope) | $R^2$ | RMSE |
| --- | --- | --- | --- | --- |
| **Stem volume** (cubic decimeter) |  |  |  |  |
| ACE1 | ACE2 | 0.64 | 0.30 | 3.17 |
| ACE1 + Mainline | ACE2 | 0.59 | 0.49 | 1.17 |
| ACE1 + ACE2 | Mainline | 0.32 | 0.11 | 2.00 |
| **Stem straightness** ( 1 to 6 scale) |  |  |  |  |
| ACE1 | ACE2 | 0.66 | 0.45 | 0.20 |
| ACE1 + Mainline | ACE2 | 0.72 | 0.49 | 0.22 |
| ACE1 + ACE2 | Mainline | 0.52 | 0.14 | 0.30 |

**Figure 3.**
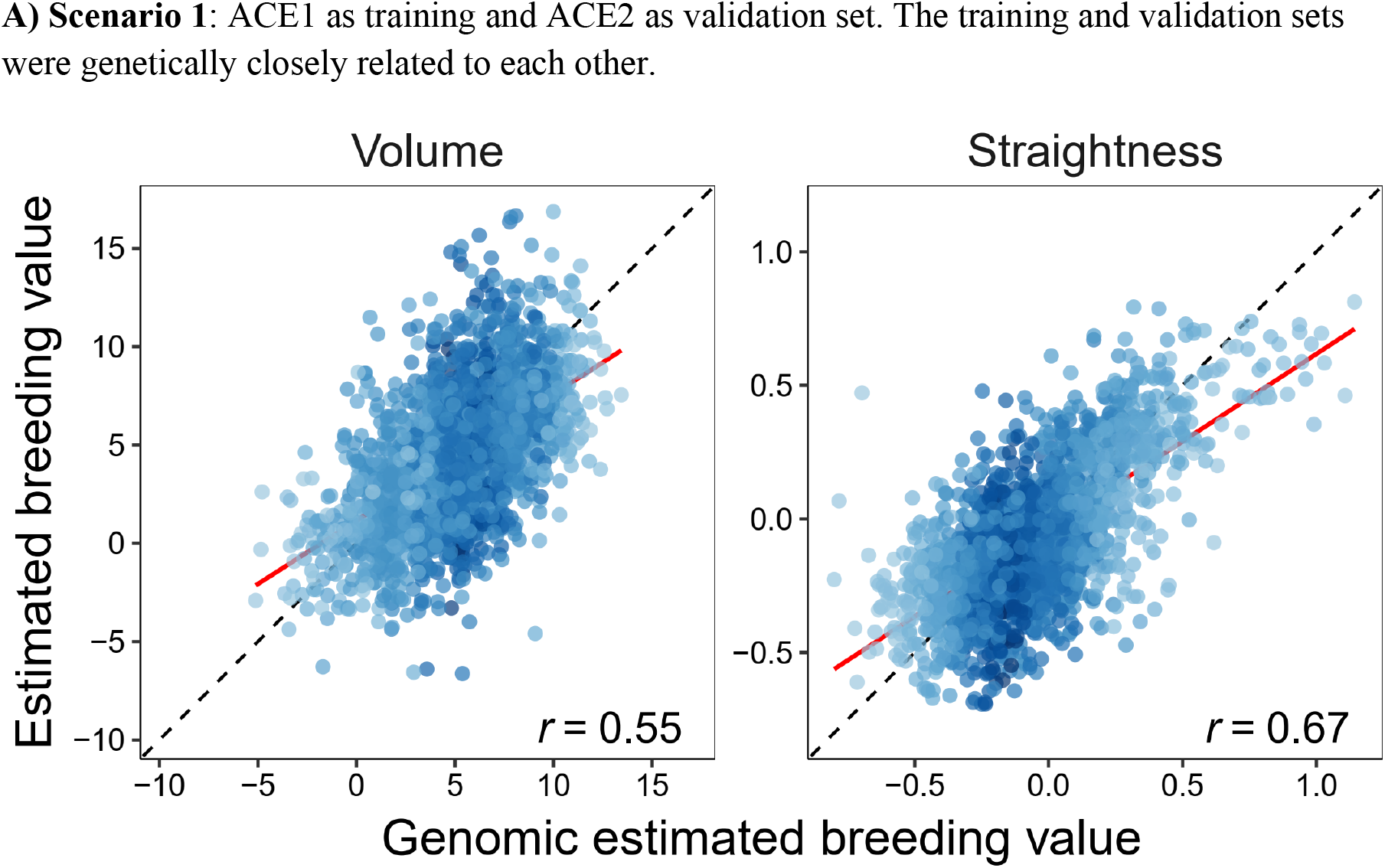
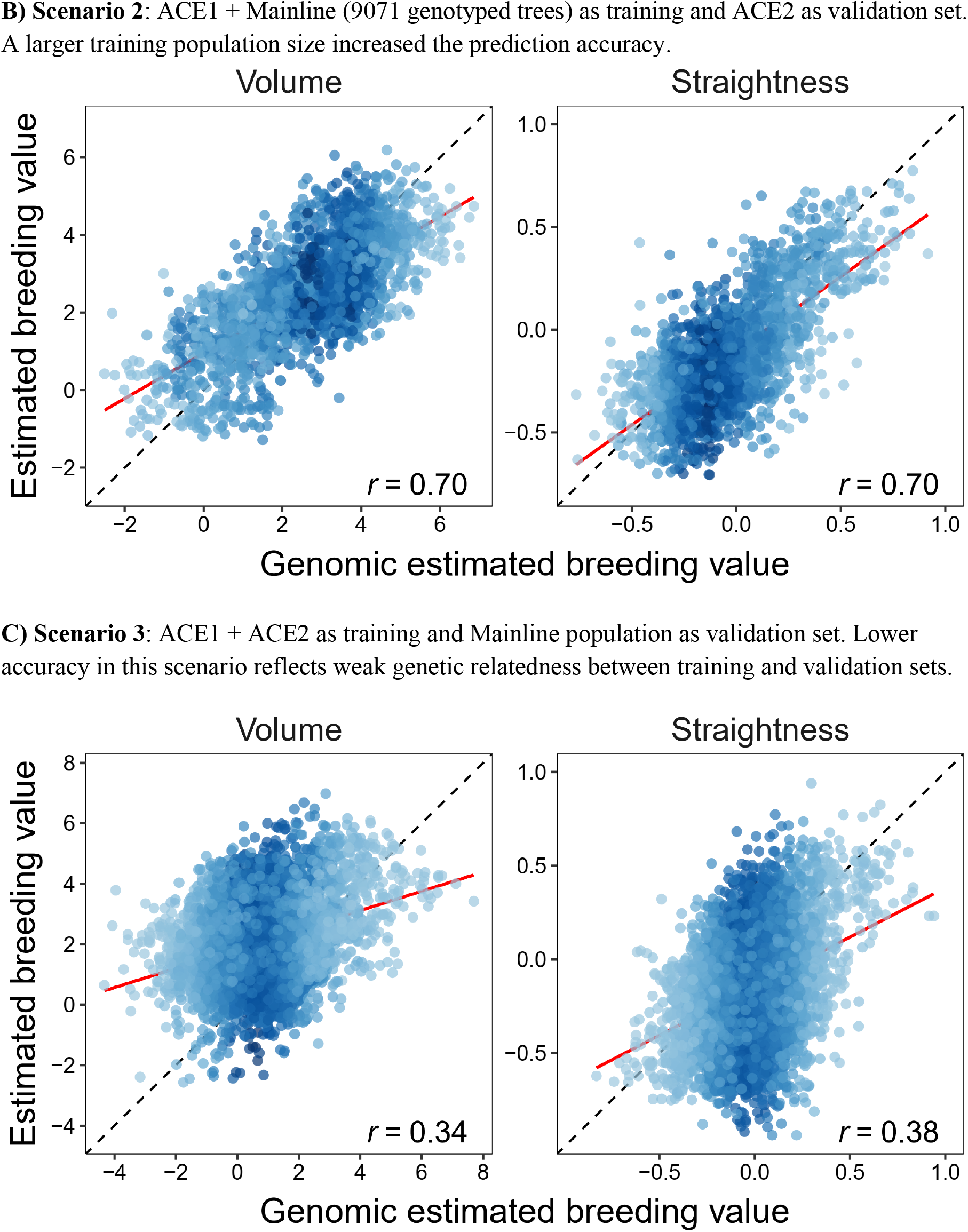
Relationships between genomic estimated breeding values (GEBVs) and pedigree-based estimated breeding values (EBVs) for stem volume and stem straightness under three validation scenarios. The dashed black line indicates the 1:1 relationship, and the solid red line shows the fitted regression. Correlation coefficients (r) represent prediction accuracy for each scenario. **A) Scenario 1**: ACE1 as training and ACE2 as validation set. The training and validation sets were genetically closely related to each other. **B) Scenario 2**: ACE1 + Mainline (9071 genotyped trees) as training and ACE2 as validation set. A larger training population size increased the prediction accuracy. **C) Scenario 3**: ACE1 + ACE2 as training and Mainline population as validation set. Lower accuracy in this scenario reflects weak genetic relatedness between training and validation sets.

In the second scenario, ACE1 and the Mainline population were combined to form a larger training set, with ACE2 retained for validation. The increase in training population size substantially improved prediction accuracy for both traits (*r* = 0.70) (**Figure 3B**). The GEBV inflation persisted for stem volume (*b*_1_ = 0.59) but reduced for straightness (*b*_1_ = 0.72) (**Table 3**). For both traits, GEBV explained 49% of the variation in EBV (*R*^2^ = 0.49).

In the third scenario, the combined ACE population (ACE1 + ACE2) was used for training, and the Mainline population served as the validation set by masking phenotypes of all genotyped individuals. Prediction accuracy declined markedly for stem volume (*r* = 0.34) and stem straightness (*r* = 0.38), accompanied by further reductions in regression slopes, indicating increased inflation of GEBVs (**Figure 3C**). This decline reflects the lower average genetic relatedness between ACE and Mainline families (**Table S1**).

Prediction accuracy was strongly associated with mean genetic relatedness between training and validation sets when using the ACE population to predict GEBVs for the Mainline population (**Figure 4**). The correlation between mean genetic relatedness and prediction accuracy was 0.97 for stem volume and 0.92 for stem straightness. This finding highlights the critical role of genetic connectivity for reliable genomic prediction.

**Figure 4.**
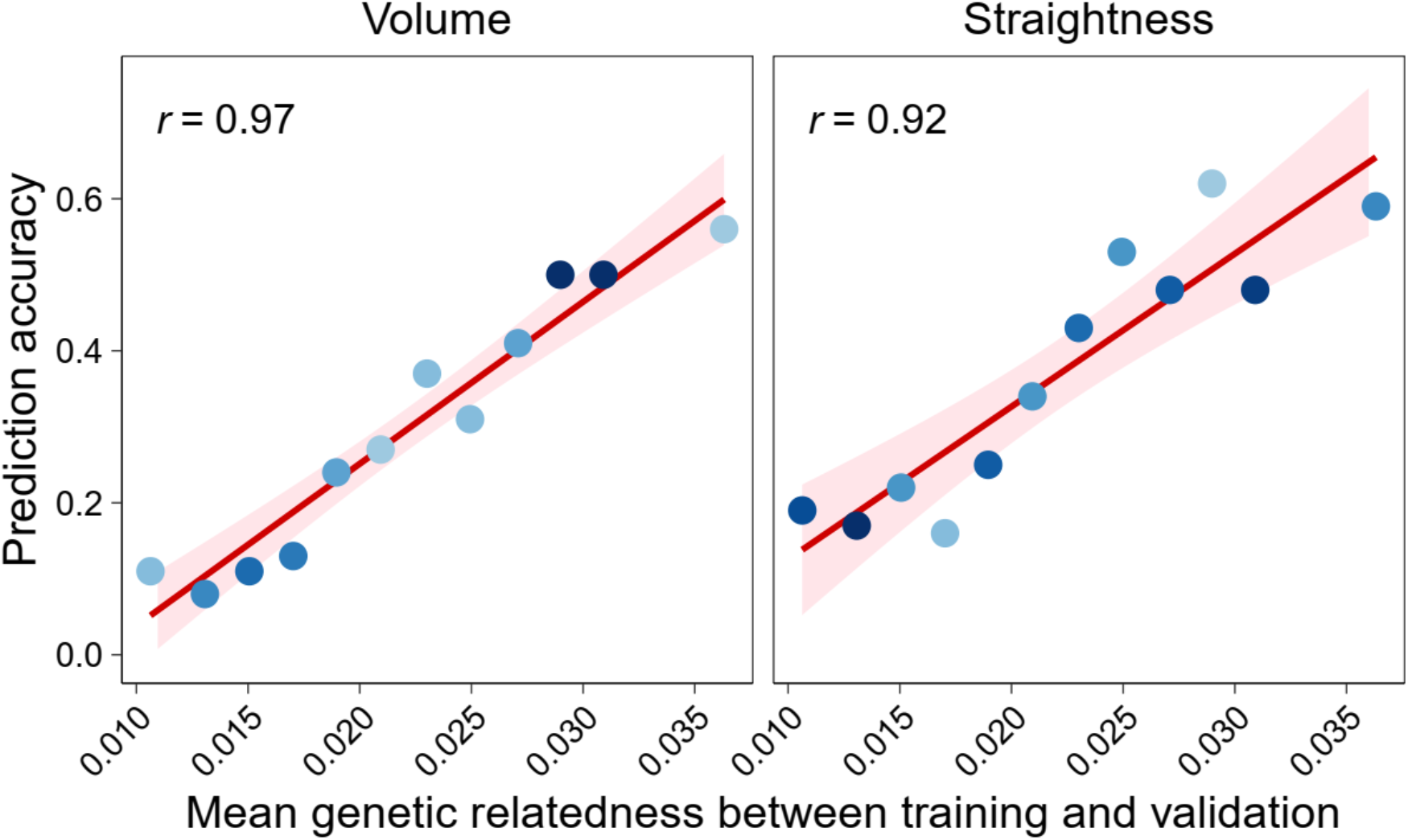
Relationship between mean genomic relatedness of training and validation populations and prediction accuracy of genomic estimated breeding values (GEBVs) for stem volume and stem straightness. Each point represents a validation subset from the Coastal Mainline population, with the Atlantic Coastal Elite (ACE) population used as the training set. Prediction accuracy increased with greater genetic relatedness between training and validation populations.

### Scaling genetic relationships

Scaling the genomic relationship to improve compatibility with the pedigree relationship using a weight factor (λ) in the construction of **H**^**−1**^ had consistent positive effects on genetic parameter estimates and genomic prediction performance in the ACE population. As λ decreased, thereby increasing the relative contribution of pedigree relationships, heritability estimates increased, prediction accuracy improved, and bias in genomic estimated breeding values (GEBVs) was reduced for both traits (**Table 4**).

**Table 4.**
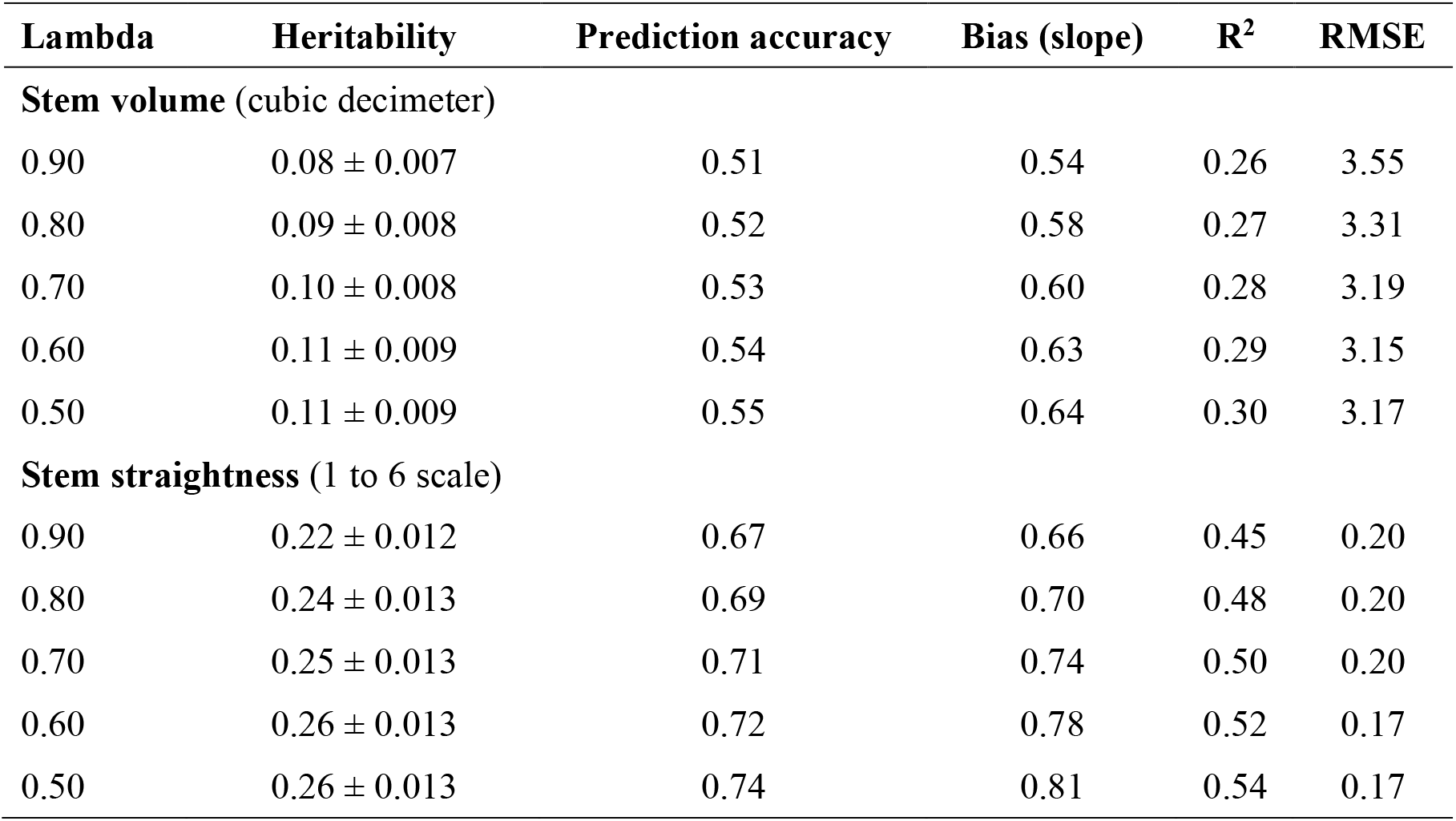
Effect of the scaling factor (*λ*) applied to marker- and pedigree-based genetic relationships in single-step genomic best linear unbiased prediction (ssGBLUP) on estimates of narrow-sense heritability and genomic prediction validation statistics for stem volume and stem straightness in the Atlantic Coastal Elite (ACE) population. Results are shown for Scenario 1, with ACE1 used as the training population and ACE2 as the validation population. Validation statistics include prediction accuracy, regression slope (calibration), coefficient of determination (*R*^2^), and root mean squared error (RMSE).

For stem volume, reducing λ (more weight on pedigree information) led to higher heritability and modest but consistent gains in prediction accuracy (0.51 to 0.55), accompanied by regression slopes closer to unity and improved overall model fit. Improvements tended to plateau at intermediate λ values, indicating diminishing returns from further increasing the weight on pedigree information. Collectively, these results support the use of λ = 0.50 in the construction of **H**^**−1**^ to improve estimation of GEBVs for stem volume in ssGBLUP model.

For stem straightness, the response to scaling was stronger. Lower λ values (more weight on pedigree) resulted in clear increases in prediction accuracy and substantial reductions in GEBV inflation, along with improved model fit. For example, heritability increased from 0.22 to 0.26, while the prediction accuracy increased from 0.67 to 0.74 when the lambda decreased from 0.90 to 0.50. Compared with stem volume, genetic parameter estimates for straightness were less sensitive to scaling, consistent with its higher heritability.

## Discussions

This study validates the practical utility of genomic selection (GS) across generations in loblolly pine, an economically important forest tree species in the southern United States. After breeding, recombination can erode linkage disequilibrium between markers and causal loci; however, the prediction accuracies observed here were comparable to those reported in early GS studies based on cross-validation (Lauer et al. 2022; Shalizi et al. 2022). Estimates of variance components and heritability were similar between pedigree-based and marker-based models, particularly after genomic relationships were scaled to be compatible with pedigree expectations. Genetic gain across generations estimated from GS was comparable to that from conventional selection. If the breeding cycle of loblolly pine shortened by approximately four years, GS is expected to increase genetic gain per year by about 50%.

We identified three major factors affecting GS prediction accuracy: genetic relatedness between training and validation populations, training population size, and scaling of genomic relationship matrices. In livestock, GS models trained within one breed often perform poorly in unrelated breeds because predictive ability depends on tracking recent shared haplotypes (Lund et al. 2014; Wiggans et al. 2017). GS studies in forest trees anticipated the importance of relatedness using cross-validation approaches (Isik 2022; Shalizi et al. 2022; Lauer et al. 2022), but true across-generation validation studies are only now emerging (Simiqueli et al. 2023). In our study, increasing mean relatedness between training and validation sets resulted in a linear increase in prediction accuracy from approximately 0.10 to 0.60, with no indication of a plateau. These results highlight the importance of investing in training populations that are well connected to the target breeding population.

Increasing the training population size from approximately 3,000 to 9,000 genotyped trees substantially improved prediction accuracy for stem volume, a trait with modest heritability (h^2^ ≈ 0.1), and resulted in a smaller improvement for stem straightness, which had higher heritability (h^2^ ≈ 0.2). Larger training populations provide more information to estimate additive genetic effects from realized relationships, particularly within large full-sib families where markers capture Mendelian sampling variance (Hayes et al. 2009). The optimal number of genotyped individuals depends on population diversity and effective population size, both of which are typically large in forest tree breeding programs.

Single-step genomic best linear unbiased prediction represents the most practical framework for implementing GS in operational tree breeding programs. This approach is now standard in livestock breeding, where the theory underlying the scaling of genomic and pedigree relationships is well established (VanRaden 2008; Christensen and Lund 2010; Bermann et al. 2022). Compared with two-stage approaches, ssGBLUP is simpler, accounts for selection, and can reduce bias in genomic estimated breeding values, provided that relationship matrices are properly scaled (Vitezica et al. 2011; Legarra et al. 2014). Despite its importance, appropriate scaling of genomic relationships has received limited attention in forest trees. Our results show that scaling can have a substantial, trait-dependent impact on variance components and prediction accuracy.

We consistently observed inflation (regression coefficients < 1) when EBVs of genotyped individuals were regressed on their GEBVs, even after scaling genomic and pedigree relationship matrices and accounting for residual polygenic effects (Christensen and Lund 2010). Inflated predictions were observed in a hybrid eucalyptus population, although those analyses were limited to a single generation and smaller population sizes (Cappa et al. 2019). In dairy cattle, bias in GEBVs of test bulls was reduced when genotyped bulls without progeny records were excluded (Koivula et al. 2018). Inflation of GEBVs is generally attributed to incompatible scaling between genomic and pedigree relationships, overweighting of genomic information, incomplete capture of additive genetic variance by markers, and limited relatedness between training and validation populations (VanRaden 2008; Christensen and Lund 2010; Misztal et al. 2009; Vitezica et al. 2011). In contrast, deflation observed in U.S. dairy cattle has been linked to incomplete pedigree information and strong selection pressure (Misztal and Legarra 2017).

### Implications for loblolly pine breeding

This study provides the first evaluation of genomic prediction across generations in a conifer tree species and highlights the importance of developing large, genetically connected training populations supported by reliable pedigrees and high-quality phenotypes. With a well-developed training population, GS can provide sufficient prediction accuracy to support recurrent genomic selection of loblolly pine. In the North Carolina State University Cooperative Tree Improvement Program, development of a GS training population began in 2019 with genotyping approximately 3,000 trees in the elite population described in this study, followed by targeted genotyping of the mainline population (fourth-cycle progeny tests) to reach a total of approximately 9,000 genotyped trees. The mainline population consisted of over 55,000 progeny test trees, and genotyping focused on families contributing fifth-cycle selections and closely related families, and trees with breeding values with low standard errors, typically from uniform test sites with excellent silviculture and high-quality phenotypes. This training population is expected to provide adequate prediction accuracy to support sixth-cycle recurrent GS.

The current GS strategy envisions genotyping approximately 3,000 full-sibling seedlings per year from about 100 controlled crosses beginning in 2026, with roughly 3% selected as parents for breeding in the sixth cycle. All genotyped seedlings will be established in replicated field trials to generate high-quality phenotypic data to retrain models for continued improvement of genomic predictions. Some programs that employ clonal forestry have suggested using GS to discard seedlings from clonal testing, effectively increasing the selection intensity or reducing the testing load (Balocchi et al. 2023; Grattapaglia 2022). In our program, we expect retaining genotyped individuals for later phenotyping as continued validation and model retraining to be a better investment.

Under conventional breeding, a full selection cycle, from progeny testing and selection through grafting, breeding, seed harvest, and establishment of new tests, requires approximately 12 years (Isik and McKeand 2019). In contrast, a recurrent GS strategy has the potential to reduce the breeding cycle by approximately four years, increasing the annual rate of genetic gain up to 50%. However, several biological and logistical challenges remain for full implementation of GS. In conventional breeding, cuttings from selected trees are typically collected at age five or six years (since seed) and grafted into the crowns of mature rootstocks to induce production of strobili for breeding. Under a recurrent GS strategy, cuttings would be collected from seedlings at one or two years of age. It remains unclear whether grafts from such young selections will produce strobili as timely as those from older trees. Further, scion size limitations may require an additional year of growth in field settings. Consequently, the anticipated four-year reduction in breeding cycle length from GS assumes onset of flowering similar to conventional selections, an assumption that is currently being empirically tested through grafting and flower induction treatments.

### Conclusions

This study demonstrates that genomic selection can effectively bypass progeny testing in loblolly pine breeding programs, providing a clear pathway to increased rates of genetic gain. Prediction accuracy was strongly influenced by training population size, genetic relatedness between training and validation populations, and the scaling of genomic and pedigree relationship matrices. These results emphasize the importance of well-developed training populations and consistent statistical frameworks. Building on these findings, plans are underway to implement genomic selection routinely in loblolly pine breeding programs beginning in 2026.

## Data availability

Data statistics are provided in Table 1. Approximate locations of field test sites are provided in Figure 1a. Pedigree structure of the populations is provided in Figure 1b. A heatmap of genetic relationships is provided in Figure S1. Data sets and the descriptions of data sets will be available once the manuscript goes through a review process. Coded data will be deposited at Zenodo.

