## Supplemental Table S1 and Figure S1 for "Genomic selection validated across two generations of loblolly pine breeding"

### Supplemental material

**Table S1.** Cross-validation scenarios involving two generations of the Atlantic Coastal Elite (ACE) population and the Mainline population of loblolly pine. Training and validation set sizes are shown for each scenario, along with mean genetic relationship between training and validation sets based on pedigree ($\bar{\mathbf{A}}$) and genomics ($\bar{\mathbf{G}}$).

| Training set | Size | Validation | Size | Genetic relationships |
|---|---|---|---|---|
| ACE1 | 2931 | ACE2 | 3639 | $\bar{\mathbf{A}}$ = 0.056, $\bar{\mathbf{G}}$ = 0.036 |
| ACE1 + Mainline | 9071 | ACE2 | 3639 | $\bar{\mathbf{A}}$ = 0.033, $\bar{\mathbf{G}}$ = 0.024 |
| ACE1 + ACE2 | 6570 | Mainline | 6140 | $\bar{\mathbf{A}}$ = 0.023, $\bar{\mathbf{G}}$ = 0.021 |

**Figure S1.** Expected additive genetic relationships based on pedigree (upper triangle) and genomic relationships (lower triangle), within and among ACE1 (n=58), ACE2 (n=67), and Mainline (n=483) families (left) and a zoomed view of relationships within and between ACE1 and ACE2 (right). The small blocks along the diagonal represent within-family genetic relationships. The relationships between families in the ACE2 population appear more granular than in ACE1, reflecting the selective breeding of a subset of ACE1 trees to create ACE2.

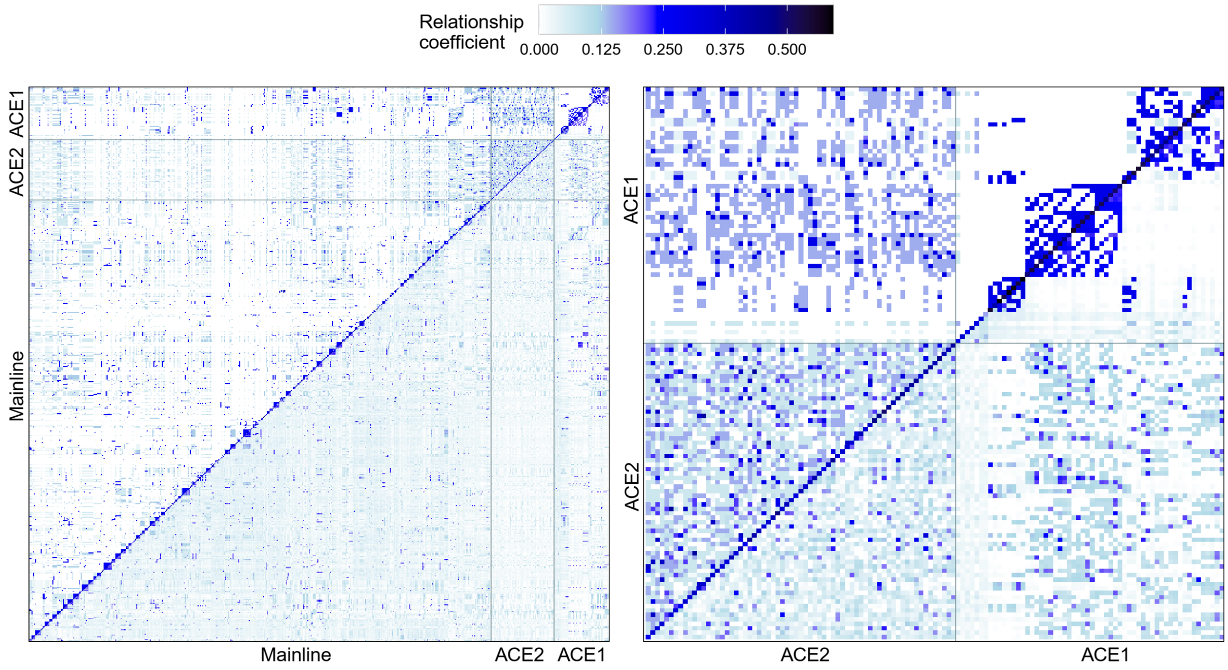
